# A second hybrid-binding domain modulates the activity of *Drosophila* ribonuclease H1

**DOI:** 10.1101/2020.03.26.010645

**Authors:** Jose M. González de Cózar, Maria Carretero-Junquera, Grzegorz L. Ciesielski, Sini M. Miettinen, Markku Varjosalo, Laurie S. Kaguni, Eric Dufour, Howard T. Jacobs

## Abstract

In eukaryotes, ribonuclease H1 (RNase H1) is involved in the processing and removal of RNA/DNA hybrids in both nuclear and mitochondrial DNA. The enzyme comprises a C-terminal catalytic domain and an N-terminal hybrid-binding domain (HBD), separated by a linker of variable length, which in *Drosophila melanogaster* (*Dm*) is exceptionally long, 115 amino acids. Molecular modeling predicted this extended linker to fold into a structure similar to the conserved HBD. We measured catalytic activity and substrate binding by EMSA and biolayer interferometry, using a deletion series of protein variants. Both the catalytic domain and the conserved HBD were required for high-affinity binding to heteroduplex substrates, whilst loss of the novel HBD led to a ∼90% drop in K[cat] with a decreased K[M], and a large increase in the stability of the RNA/DNA hybrid-enzyme complex. The findings support a bipartite binding model for the enzyme, whereby the second HBD facilitates dissociation of the active site from the product, allowing for processivity. We used shotgun proteomics to identify protein partners of the enzyme involved in mediating these effects. Single-stranded DNA-binding proteins from both the nuclear and mitochondrial compartments, respectively RpA-70 and mtSSB, were prominently detected by this method. However, we were not able to document direct interactions between mtSSB and *Dm* RNase H1 when co-overexpressed in S2 cells, or functional interactions *in vitro*. Further studies are needed to determine the exact reaction mechanism of *Dm* RNase H1, the nature of its interaction with mtSSB and the role of the second HBD in both.

## INTRODUCTION

In eukaryotes, ribonuclease H1 (RNase H1) is present both in mitochondria and in the nucleus. The enzyme digests the RNA strand of RNA/DNA heteroduplexes longer than 4 bp, and has been demonstrated to influence diverse DNA transactions in one or both cellular compartments, including DNA replication, transcription, recombination, repair and telomere maintenance. The targeting of the enzyme to two cellular compartments complicates functional studies. Therefore, our understanding of its biology has instead relied heavily on biochemical analysis.

Eukaryotic RNase H1 presents a conserved domain organization, with a nucleic-acid binding motif, denoted the hybrid-binding domain (HBD), also found in some bacteria, such as *Bacillus halodurans* (3) towards the N-terminus, a catalytic (RNase H) domain located near the C-terminus and an extended linker between these two domains. In human RNase H1, the HBD confers tight binding to RNA/DNA hybrid, but also interacts weakly with dsRNA and even more weakly with dsDNA, and portions of its structure that interact with RNA and with DNA have been mapped precisely (4). The physiological role of the HBD remains unknown, but it has been suggested to promote dimerization, conferring processivity to the enzyme (5), as well as interactions with other proteins (6). Since the first RNase H structure from bacteria was elucidated in 1990 (7, 8), a conserved three-dimensional organization has been observed in homologs of the enzyme from eukaryotes (9), prokaryotes (10, 11) and viruses (12-14). This specific structure involves 4 residues in close proximity, the DDED motif (15), serving as the catalytic core of the enzyme (see Fig. S1F). The catalytic domain of human RNase H1 has been studied extensively with respect to its binding affinity (16), cofactor requirements and structure (9). Among eukaryotes, the extended linker that connects the HBD and RNase H (catalytic) domains is the least phylogenetically conserved portion of the protein, both in length and primary sequence (Fig. 1; Ref. 17). The human linker spans 64 amino acids, whereas it is 48 residues long in *C. elegans* and 115 in *Drosophila melanogaster*. The current view is that the extended linker of the human enzyme confers flexibility to the RNase H domain whereas the HBD remains bound to the substrate (18); however, the variability of the linker among species remains unexplained.

**Figure 1.**
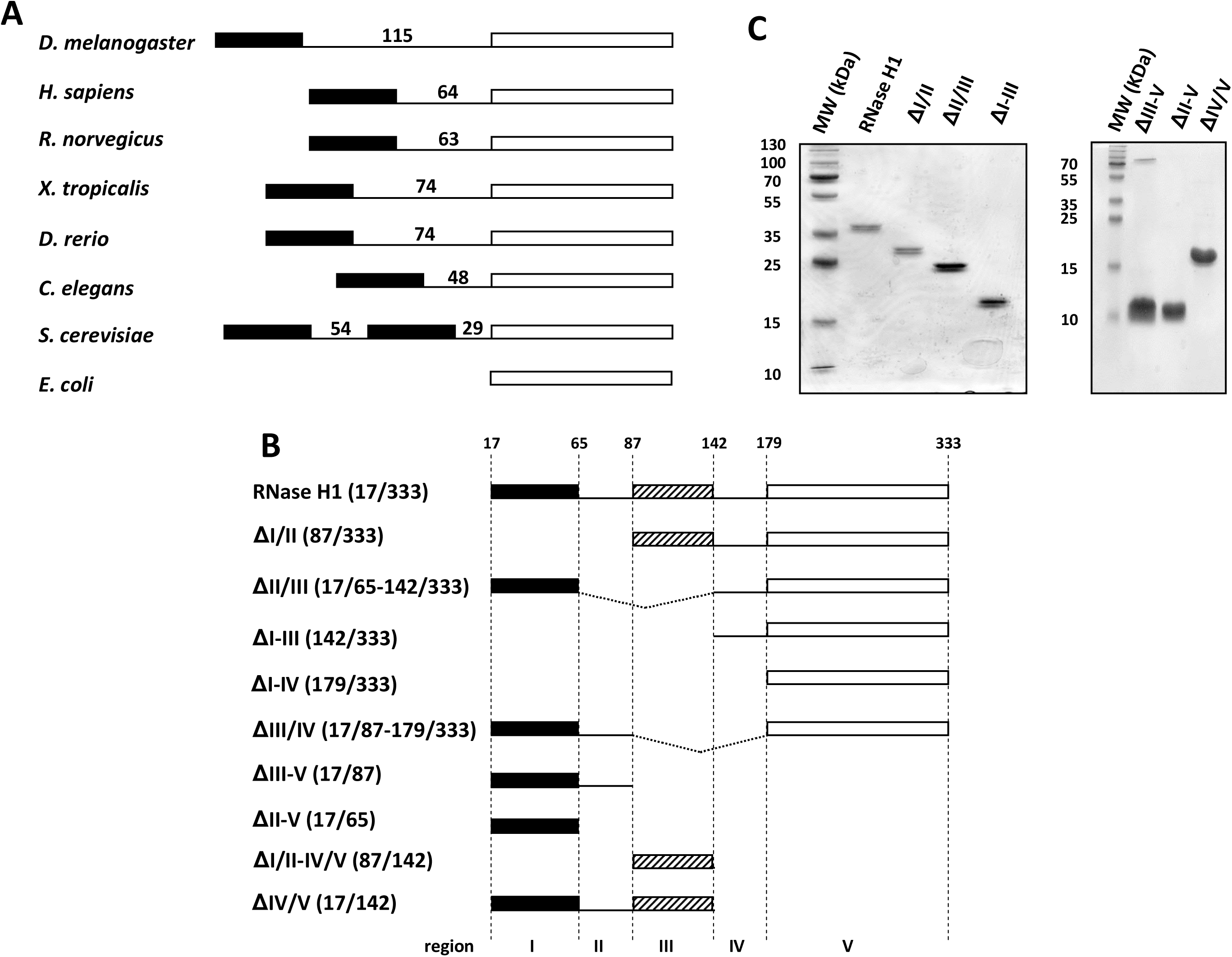
*Drosophila melanogaster* (*Dm*) RNase H1 protein sequence organization and structure. (A) Schematic representation of RNase H1 enzyme among different species (*Drosophila melanogaster, Homo sapiens, Rattus norvegicus, Xenopus tropicalis, Danio rerio, Caenorhabditis elegans, Saccharomyces cerevisiae* and *Escherichia coli*). Black box represent the conserved HBD and the empty box the catalytic domain. Length of the linker region (in amino acids) for each species is shown. Note that *S. cerevisiae* has two HBDs and two linkers (41). (B) Schematic representation of different *Dm* RNase H1 protein variants created in the study: RNase H1 (Rnh1, NCBI AAF59170.1 amino acids 17-333), ΔI/II (87-333), ΔII/III (17-65+142-333), ΔI-III (142-333), ΔI-IV (179-333), ΔIII/IV (17-87+179-333), ΔII-V (17-65), ΔIII-V (17-87), ΔI/II-IV/V (87-137) and ΔIV/V (17-142). The whole RNase H1 protein sequence has been into 5 regions (assumed to represent protein domains) delimited by amino acids 17, 65, 87, 142, 179 and 333. Black box represents the conserved HBD, the hatched box the second, predicted HBD and the empty box the catalytic domain. (C) Purified proteins, as indicated, separated by SDS-PAGE and Coomassie staining, alongside molecular mass (MW) markers in kDa.

The *ribonuclease H1* (*rnh1*) gene is necessary for pupal development in *Drosophila melanogaster* (19). *Rnaseh1* knockout mice also exhibit developmental lethality, with a decreased copy number of mitochondrial DNA (mtDNA), suggesting an essential role of RNase H1 in mtDNA maintenance (20). Several point mutations in the human *RNASEH1* gene are associated with progressive external ophthalmoplegia (21-23), a syndromic disorder associated with mutations in various mtDNA maintenance proteins, such as the mitochondrial DNA polymerase gamma (Polγ; Ref. 24) or mitochondrial transcription factor A (TFAM; Ref. 25). Both in mammals and *Drosophila*, RNase H1 has been shown to be simultaneously targeted to mitochondria and the nucleus (1, 2), and has been proposed to be involved in various processes requiring RNA/DNA turnover. In mitochondria, RNase H1 has been implicated in mtDNA replication initiation and segregation (2, 26, 27), as well in mitochondrial RNA processing (2, 28, 29) and RNA/DNA removal (30). In the nucleus, it has been proposed to be essential for some types of DNA repair (31, 32), removal of persistent heteroduplex (33, 34) and telomere maintenance (35).

It remains unclear whether the lethality associated with *rnh1* deletion in *Drosophila* (19) is due to a requirement for its function in the nucleus, in mitochondria or both. Recently, we described the effects of *Drosophila rnh1* downregulation, which triggers decreased lifespan, mitochondrial dysfunction and mtDNA depletion (2). This phenotype was associated with abnormalities in mtDNA replication, notably at the origin and in regions where the transcription machinery progresses in the opposite direction to that of DNA replication (2).

In the present study, in view of the preliminary findings of previous investigators (36), we set out to determine the biochemical properties of the different domains of *Dm* RNase H1, notably that of the extended linker, via functional studies of a deletion series *in vitro*, and by screening for interacting proteins. This leads us to propose a new conceptual mechanism facilitating the processivity of the enzyme, as well as an interaction with mitochondrial single-stranded DNA-binding protein (mtSSB) that appears to be robust but indirect

## RESULTS

### Structure modeling of *Drosophila* RNase H1

*Dm* RNase H1 was previously proposed to comprise three structural elements (36): an N-terminal HBD domain similar to a region of caulimovirus ORF VI protein (37), an RNase H catalytic domain located towards the C-terminus and a 115 amino acid extended linker that connects these domains. Although the extended linker is longer, the overall architecture is similar to that of RNase H1 from other eukaryotes (Fig. 1A; Ref.18). Structure modeling of the *Drosophila* sequence revealed conserved and potentially novel features of the enzyme. Two of the four amino acids of the human HBD required for hybrid binding (4) are conserved, namely Y29 and K60, contacting the DNA backbone, equivalent to Y19 and K50 in the fly enzyme (Fig. S1A). The main-chain amides of human R52 and A55 form hydrogen bonds with 2′-OH groups from two consecutive ribonucleotides. These amino acids are replaced in *Drosophila* by G42 and N45. The DEDD motif at the catalytic core (15) is also present in *Drosophila* at positions D187, E228, D252 and D318 (Fig. S1D, S1F) as are two of the three amino acids of the catalytic domain involved in hybrid binding (9) in human, i.e. N151 and N182 (N193 and N224 in *Drosophila*), whilst the third, human Q183 is represented in *Drosophila* as N225. Protein structure modeling software predicted that a portion of the extended linker could fold as an additional HBD (amino acids 87-142, Fig. S1B, S1C). The conserved N-terminal HBD (amino acids 17-64) shows 42% sequence identity with human RNase H1, with a QMEAN (estimated value of the geometric properties of the model) of 0.55 and a GMQE (global model quality estimation: being a quality estimation that merges properties of the sequence-template alignment and the template search method) of 0.18. The second, predicted HBD has only 24% sequence identity, a QMEAN of −3.32 and GMQE of 0.08 (38-40). The conserved HBD shows a similar structure as in human (4), with two α-helices and 3 antiparallel β-sheets, organized as a ββαβα structure. The second, predicted HBD is similar, but with longer loops connecting β_1_ to β_2_ and α_1_ to β_3_. The presence of an additional HBD has previously been reported in *Saccharomyces cerevisiae* (17, 41), where it stabilizes interaction with dsRNA. These observations prompted us to explore the binding properties of the different domains of the *Drosophila* enzyme.

### Biochemical characterization of *Drosophila melanogaster* RNase H1

To explore its biochemical properties, the full-length *Dm* RNase H1 protein was expressed in *E. coli*. The expressed protein consisted of 316 amino acids from the RNase H1 open-reading frame, commencing at A17, assuming post-translational removal of the N-terminal formyl-methionine by the action of methionine aminopeptidase (42), plus an additional 8 amino acids at the C-terminus from the 6xHis-tag and two further amino acids contributed by the vector, pET-26b(+). This expressed protein represents the nuclearly-targeted variant, which is very similar to the mitochondrially-targeted variant after removal of the predicted mitochondrial targeting signal. Following affinity purification and size-exclusion chromatography, it migrated on SDS-PAGE gels with an apparent molecular mass of approximately 38 kDa (Fig. 1C), close to prediction (35.1 kDa plus the C-terminal tag). We initially investigated the kinetic properties of the enzyme at 30 °C, using a blunt-ended model substrate comprising a 5′-radiolabeled 30 nt RNA oligonucleotide hybridized to a complementary 30 nt DNA oligonucleotide (Table 1: for original gels and graphical plots see Fig. S2). For comparison with the previously studied human enzyme, we reinvestigated RNase H1 from the two species at both 30 and 37 °C. At 30 °C, the human enzyme had no detectable activity, whilst at 37 °C it performed as previously reported (16), whereas the *Drosophila* enzyme showed a greatly increased K_M_ but also an approximate doubling of K_cat_ at 37 °C, compared with its properties at 30 °C. Note that 37 °C is far above the physiological temperature range for the fly enzyme *in vivo* (15-30 °C). In comparison with the human enzyme at 37 °C, the *Drosophila* enzyme at 30 °C can be regarded as considerably more active, with a much lower K_M_ and much higher K_cat_ (Table 1, Fig. S2).

**Table 1.** Kinetic parameters^a^ of RNase H1 and variants.

| Enzyme | $K_M$ (nM) | $K_{cat}$ (min<sup>-1</sup>) |
| --- | --- | --- |
| Human RNase H1 (37°C) | 41.4 ± 8.22 | 3.00 ± 1.49 |
| *Dm* RNase H1 (37°C) | 171 ± 5.7 | 27.1 ± 2.52 |
| *Dm* RNase H1 (30°C) | 8.45 ± 1.84 | 13.0 ± 3.29 |
| ΔI/II (30°C) | 4.01 ± 1.05 | 12.7 ± 7.7 |
| ΔII/III (30°C) | 3.12 ± 0.55 | 1.63 ± 1.61 |
| ΔI-III (30°C) | 9.69 ± 1.78 | 30.3 ± 8.95 |
| *Dm* RNase H1 D252N (30°C) | 0 | 0 |

<sup>a</sup>Means ± SD (n ≥ 3) for *Dm* and human enzymes and variants. All values given to 3 significant figures.

To study the binding of *Dm* RNase H1 and its derivatives to nucleic acid substrates, we generated a single point mutation (D252N) at the catalytic core, which abolished enzymatic activity (Fig. 2A). This enabled us to use an electrophoretic mobility shift assay (EMSA) to profile its nucleic acid-binding properties. A ten-fold molar excess of the protein was sufficient to generate a detectable complex with a 30 bp RNA/DNA hybrid, with a secondary mobility shift seen at higher protein/hybrid ratios (Fig. 2B). Substrate-binding kinetics were then analyzed by biolayer interferometry (BLI).

**Figure 2.**
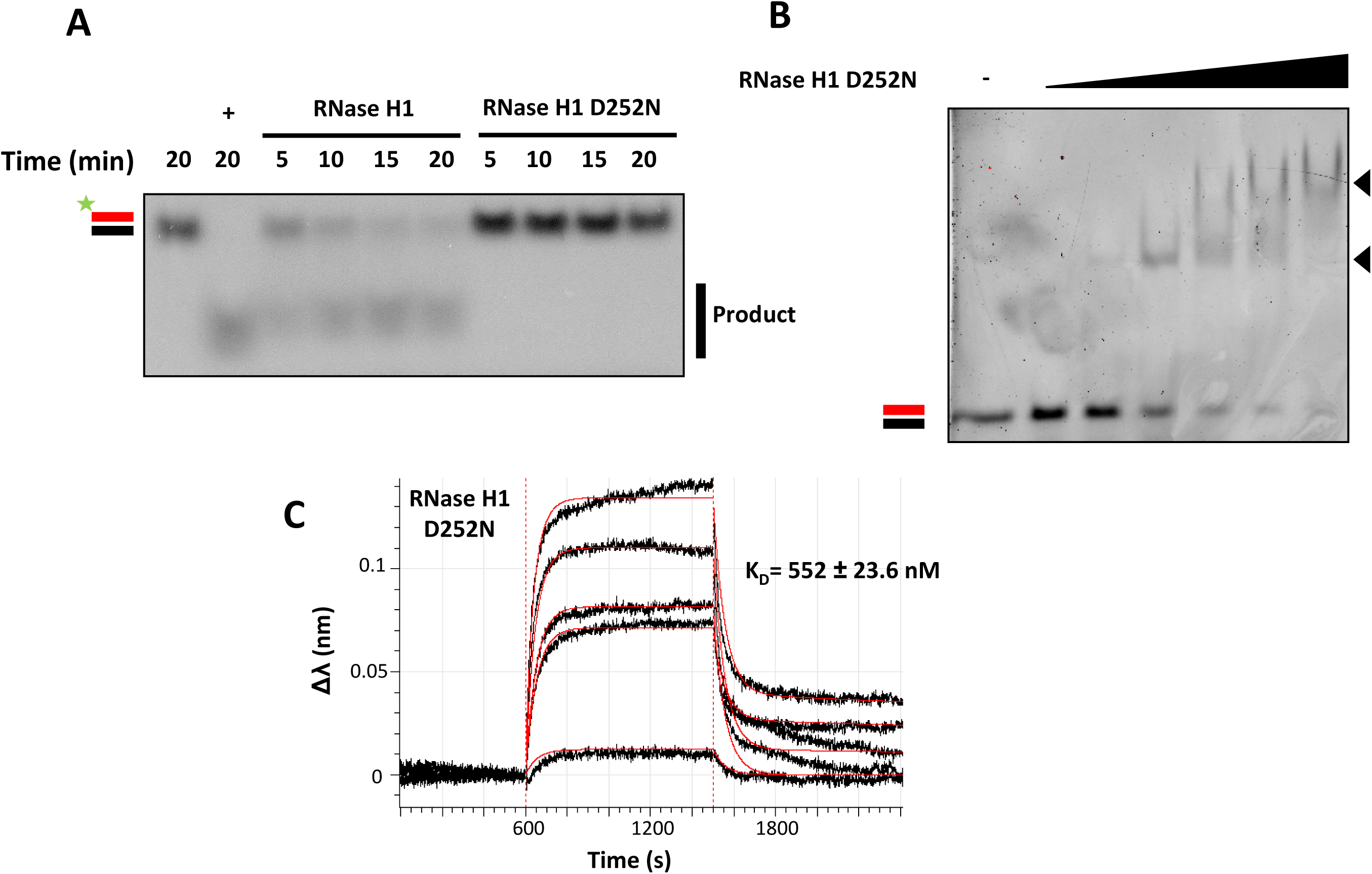
Enzymatic analysis of *Dm* RNase H1. (A) In a 10 μl reaction, 100 fmol of 30 bp 5′-radiolabeled RNA/DNA hybrid were incubated with 2 fmol of RNase H1 or RNase H1 D252N (catalytically inactive) for the indicated times (min), and separated by non-denaturing gel electrophoresis. In this and subsequent figures RNA/DNA hybrid is denoted by parallel red (RNA) and black (DNA) bars, with radiolabel indicated by the green asterisk. Positive control (+) used *E. coli* RNase H. (B) In a 20 µl reaction, 2 pmol of a 30 bp RNA/DNA hybrid were incubated with increasing concentrations of RNase H1 D252N (0, 0.1, 0.5, 1, 2.5, 5 and 10 µM) and separated by non-denaturating gel electrophoresis. Black arrowheads indicate two different protein-nucleic acid complexes. (C) BLI analysis of catalytically inactive (D252N) RNase H1 interacting with linear 30 bp RNA/DNA hybrid. Streptavidin sensors were incubated in 80 µl of 25 nM 5′ biotinylated hybrid solution. Association was measured by transferring sensor to 80 µl of different concentrations of RNase H1 D252N solution (0, 10, 20, 40, 60, 80 and 100 nM). The sensogram displays the baseline, association and dissociation steps, with experimental data shown as black lines and extrapolated data fitted to heterogeneous (2:1) binding model as red lines. See Table 2 for association/dissociation parameters.

The RNA/DNA hybrid substrate was immobilized on the sensors and the binding of the enzyme at various concentrations was analyzed in real time. The obtained association plots (Fig. 2C) were indicative of heterogeneous binding and the data were fitted well to a 2:1 binding model, wherein the enzyme binds in two steps to the substrate. The substrate-binding affinity in the first binding step was relatively low with K_D_ = 552 ± 23.6 nM. In the second binding step, the substrate-binding affinity was relatively strong, with K_D_ = 0.23 ± 0.0632 nM (Table 2). These data suggest that the first binding step may serve as the substrate recognition, whereas the second step likely serves to stabilize the enzyme on the substrate. Notably, the substrate affinity in the second step is remarkably strong, the K_D_ being ∼10-fold lower than that reported for mtSSB from *D. melanogaster* in its binding to ssDNA (2.5 nM; Ref. 43).

**Table 2.** Kinetic parameters^a^ of binding of RNase H1 variants to different substrates.

| **Nucleic acid** | **Enzyme** | **K<sub>D1</sub> (nM)** | **K<sub>D2</sub> (nM)** | **K<sub>on1</sub> (1/M\*s)\*10<sup>-3</sup>** | **K<sub>on2</sub> (1/M\*s)\*10<sup>-3</sup>** | **K<sub>off1</sub> (1/s)\*10<sup>3</sup>** | **K<sub>off2</sub> (1/s)\*10<sup>3</sup>** |
| --- | --- | --- | --- | --- | --- | --- | --- |
| **Linear hybrid** | RNase H1 | 552 ± 23.6 | 0.23 ± 0.0632 | 32.5 ± 1.39 | 490 ± 9.11 | 17.9 ± 0.0614 | 0.114 ± 0.00226 |
|  | ΔII/III | 110 ± 0.75 | 0.618 ± 0.0175 | 216 ± 1.43 | 171 ± 0.993 | 23.7 ± 0.0438 | 0.106 ± 0.00292 |
| **dsRNA** | RNase H1 | 2,170 ± 170 | 744 ± 6.21 | 202 ± 14.9 | 0.402 ± 0.0022 | 438 ± 11.7 | 0.299 ± 0.00189 |
|  | ΔII/III | 2,030 ± 30 | 934 ± 163 | 154 ± 21 | 0.658 ± 110 | 313 ± 1.93 | 0.615 ± 29.7 |
| **dsDNA** | RNase H1 | 13,500 ± 913 | 344 ± 6.55 | 35.5 ± 0.232 | 1.79 ± 0.0255 | 481 ± 8.36 | 0.618 ± 0.0746 |
|  | ΔII/III | 1,970 ± 53.8 | 116 ± 6.33 | 188 ± 4.97 | 0.629 ± 0.0070 | 371 ± 3.48 | 0.0726 ± 0.00389 |

<sup>a</sup>based on BLI, using a heterogeneous (2:1) binding model for these variants, which gave the best fit. All values given to 3 significant figures.

### HBD and catalytic domains influence RNase H1 activity and nucleic acid binding

Next we studied the properties of *Dm* RNase H1 variants bearing deletions of specific domains. For the purposes of this analysis we defined five regions of the enzyme, from N- to C-terminus (Fig. 1B), as follows: region I is the conserved HBD (amino acids 17-64), region II (amino acids 65-86) is the short linker leading up to the second, predicted HBD, region III (amino acids 87-141). Region IV is another short linker (amino acids 142-178), leading up to region V, the RNase H catalytic domain (amino acids 179-333). Variants lacking region IV but retaining region V, as well as the one comprising only region III, were insoluble and not studied further. All purified variant proteins migrated on SDS-PAGE gels approximately as predicted (Fig. 1C). The measured kinetic parameters of those variants that retained detectable catalytic activity are indicated in Table 1 (Fig. S2). In summary, deletion of regions I-III (ΔI-III) or of just the conserved HBD and its adjacent linker (ΔI/II) facilitated catalysis (Fig. 3A), though with subtly different effects on the kinetic parameters (Table 1, Fig. S2), whereas deletion of just the second predicted HBD and its upstream linker (ΔII/III) caused almost a 90 % decrease in K_cat_ at 30 °C, despite a decreased K_M_ (Table 1, Fig. 3A, S2,). In more general terms, the conserved HBD appeared to restrain catalysis, whilst the predicted second HBD prevented this repression. Both the I/II and II/III regions decreased substrate binding to the catalytic center, as inferred from lower K_M_ values in their absence. This might be a simple consequence of the presence of additional binding steps preceding the loading of the substrate to the catalytic center. In addition, the II/III region appears to support an efficient turnover rate, as its absence results in a decrease of the K_cat_ value by ∼8-fold (Table 1). A simultaneous decrease in both K_M_ and K_cat_, as observed in the absence of the II/III region (Table 1), is indicative of a decreased rate of dissociation of the enzyme-product complex. In turn, this implies that the second HBD may be relevant for effective product release. Given that the RNase has to progress along the substrate, the binding properties of domain III might be relevant to the translocation process. The lack of both HBD domains resulted in a greater than 2-fold *increase* in turnover rate (Table 1), which implies that substrate binding by the HBD domains together limits the rate of catalysis.

**Figure 3.**
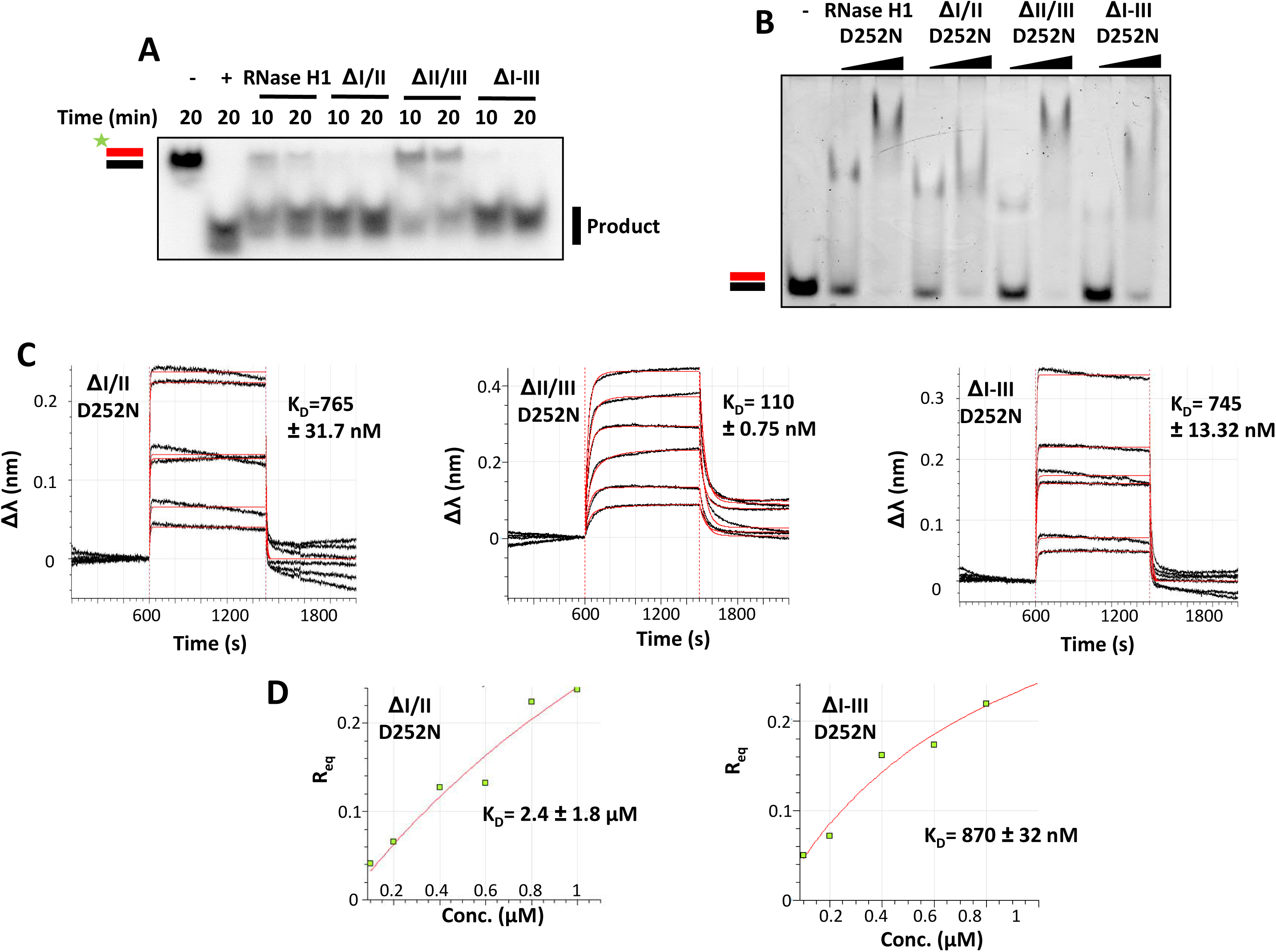
Biochemical analysis of *Dm* RNase H1 variants. (A) Enzymatic assay, (B) EMSA and (C, D) BLI, using *Dm* RNase H1 variants as indicated. Other details as for Fig. 2, except protein concentrations in (B) were 1 and 10 μM. Positive control in (A) used *E. coli* RNase H (+). For BLI analysis, sensors were incubated in increasing concentrations of RNase H1 variant proteins (ΔI/II and ΔI-III: 0, 100, 200, 400, 600, 800 and 1000 nM; ΔII/III: 0, 10, 20, 40, 60, 80 and 100 nM). The red lines of the sensograms shown in (C) are extrapolated data fitted to a monovalent (1:1) binding model (left and right sensograms, for ΔI/II and ΔI-III variants, respectively) or to a heterogeneous (2:1) binding model (central sensogram, for ΔII/III variant). See Tables 2 and 3 for association/dissociation parameters. (D) Steady-state analysis of relative equilibrium (R_eq_) plotted against protein concentration, for the variants that fitted a 1:1 binding model. Estimated dissociation constant ± standard deviation (SD) as shown.

We proceeded to test the hybrid-binding properties of these variants by introducing the D252N mutation. Analyses by EMSA (Fig. 3B) and BLI (Fig. 3C and Table 2) showed that the ability to bind RNA/DNA hybrid was retained, despite the deletion of the conserved HBD (ΔI/II D252N) or the second HBD (ΔII/III D252N) or even of both (ΔI-III D252N). However, the kinetic parameters indicated a difference in the nature of the binding. BLI data for RNase H1 variants missing the conserved HBD fitted better to a 1:1 substrate-binding model (Fig. 3 and Table 3), rather than the 2.1 model (Fig. 2, 3 and Table 2) that was more compliant with the data from full-length RNase H1 or the variant lacking only the second HBD (ΔII/III D252N). The ‘supershift’ band observed by EMSA with the full-length protein at high protein:substrate ratio was also abolished when the conserved HBD was absent (Fig. 3B).

**Table 3.**
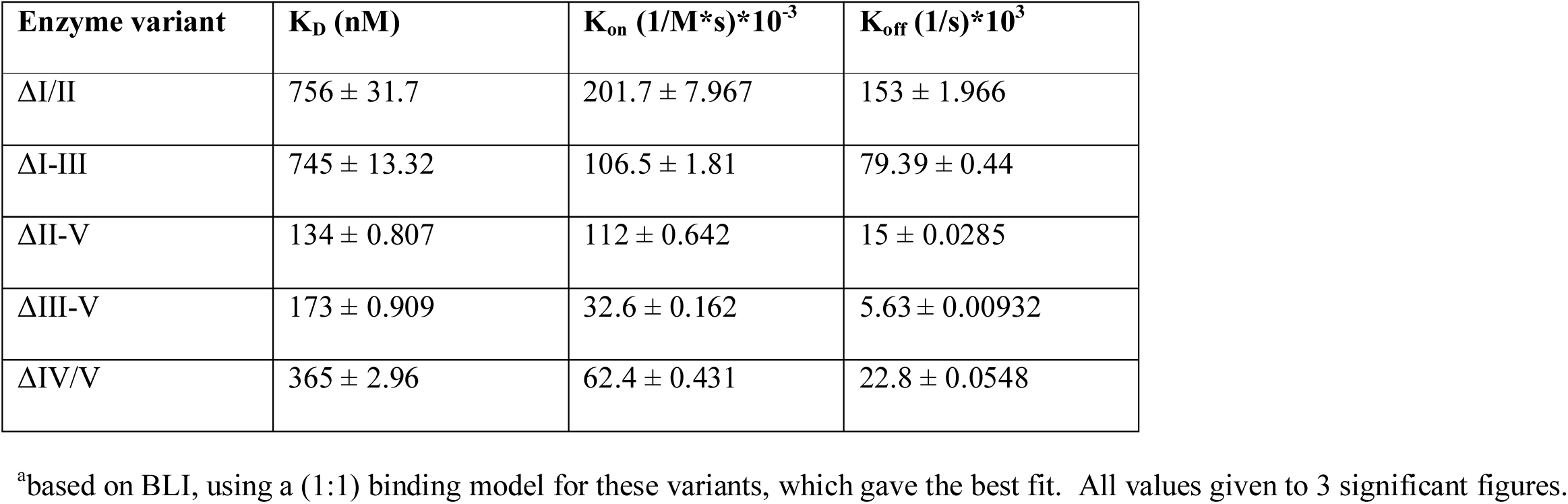
Kinetic parameters^a^ of binding to 30 bp linear hybrid, of RNase H1 variants lacking the conserved HBD or catalytic domain.

| Enzyme variant | $K_D$ (nM) | $K_{on}$ (1/M*s)*$10^{-3}$ | $K_{off}$ (1/s)*$10^3$ |
| --- | --- | --- | --- |
| $\Delta I/II$ | $756 \pm 31.7$ | $201.7 \pm 7.967$ | $153 \pm 1.966$ |
| $\Delta I-III$ | $745 \pm 13.32$ | $106.5 \pm 1.81$ | $79.39 \pm 0.44$ |
| $\Delta II-V$ | $134 \pm 0.807$ | $112 \pm 0.642$ | $15 \pm 0.0285$ |
| $\Delta III-V$ | $173 \pm 0.909$ | $32.6 \pm 0.162$ | $5.63 \pm 0.00932$ |
| $\Delta IV/V$ | $365 \pm 2.96$ | $62.4 \pm 0.431$ | $22.8 \pm 0.0548$ |

<sup>a</sup>based on BLI, using a (1:1) binding model for these variants, which gave the best fit. All values given to 3 significant figures.

Furthermore, the deletion of region I/II (conserved HBD) resulted in a ∼2-fold decrease in substrate-binding affinity (Fig. 3C, 3D and Tables 2, 3), consistent with the role of this domain in substrate binding as inferred from the lower K_M_. Conversely the lack of region II/III (the second HBD) substantially increased substrate affinity (∼5-fold) which, taken together with the decreased K_cat_, strengthens the case that it is needed for efficient substrate release and thus, most likely, for the enzyme’s translocation along the substrate. The full N-terminal region (I-III) truncation behaved in a similar manner to the variant lacking only the conserved HBD (Table 3), implying that the second HBD only functions in co-operation with the first.

We next considered the broader nucleic-acid binding properties and substrate preferences of the enzyme. EMSA analysis using the D252N-substituted variants revealed that the full-length protein, as well as deletion constructs lacking the second HBD (ΔII/III), were able to bind both dsDNA (Fig. 4A) and dsRNA (Fig. 4B), although this binding was weakened substantially when the conserved HBD (ΔI/II) or both HBDs (ΔI-III) were deleted (Fig. 4A, 4B). An R-loop substrate was bound by all of these constructs, including the catalytic domain alone, together with the preceding short linker (ΔI-III; Fig. 4C), and in each case a supershift was observed at high protein concentration (Fig. 4C). BLI (Table 2) confirmed these findings, although the deletion of the second, predicted HBD increased the binding affinity for dsDNA, but not dsRNA (Fig. S3, S4). The affinity of all of the tested variants for RNA/DNA hybrid was at least 1-2 orders of magnitude greater than for dsRNA or dsDNA (Table 2). Neither the full-length D252N-substituted protein, nor any of the variants tested, had any detectable affinity for ssRNA or ssDNA (Fig. S5) nor did the equivalent variants without the D252N substitution show any detectable nuclease activity against dsDNA, (Fig. 4D), dsRNA (Fig. 4E), ssDNA (Fig. 4F) or ssRNA (Fig. 4G). Removal of the second HBD, but not the conserved or both HBDs, had a negative effect on catalytic activity using the R-loop substrate (Fig. 4H), although there was no apparent stimulation of activity by removal of the conserved HBD, as was seen with the linear RNA/DNA substrate (Fig. 3A). Another difference between the linear hybrid and R-loop substrates was that the constructs lacking the conserved HBD still produced an EMSA supershift using the latter substrate (Figure 4C).

**Figure 4.**
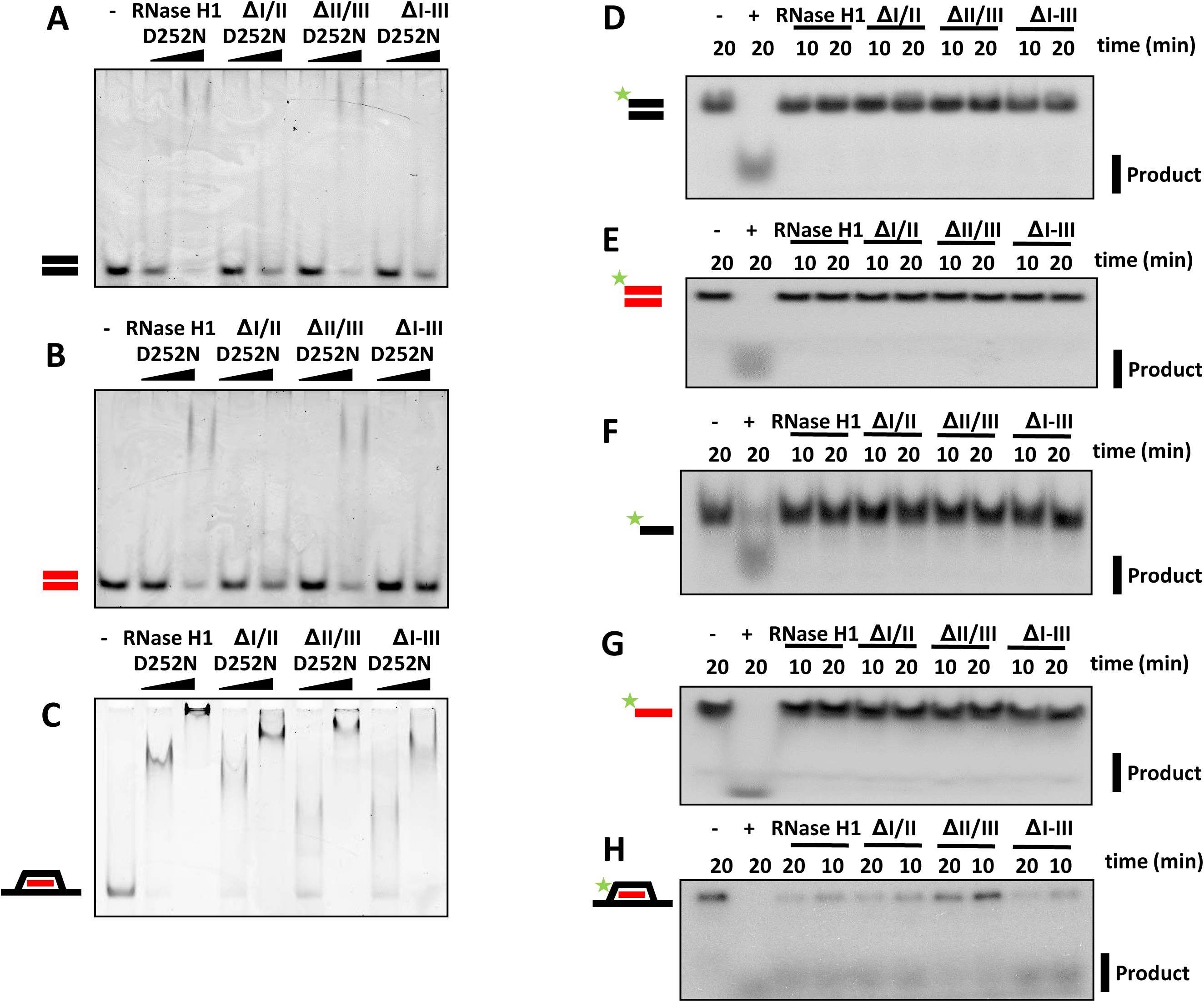
*Dm* RNase H1 binds but does not degrade non-hybrid double-stranded nucleic acid. (A, B, C) EMSA and (D-H) nuclease assays using *Dm* RNase H1 variants and substrates as indicated, according to the nomenclature of Fig. 2. Protein concentrations in (A, B, C) as for Fig. 3, other details as for Fig. 2. Positive control (+) in (D, F) DNase I, (E, G) RNase A and (H) *E. coli* RNase H.

Finally, we studied the binding properties of D252N-substituted variants ΔII-V, ΔIII-V and ΔIV/V, lacking the catalytic domain. In EMSA, all of these variants bound RNA/DNA hybrid and showed a supershift at high protein concentration (Fig. 5A). However, none of them bound dsDNA (Fig. 5B, Fig. S6A) or dsRNA (Fig. 5C, Fig. S6B). Quantitative analysis by BLI (Fig. 5D, 5E, Table 3) showed that these variants all bound hybrid more tightly than the full-length protein, with the conserved HBD alone (ΔII-V) giving the strongest binding. Overall, these findings confirm that the conserved HBD confers tight binding to hybrid, whilst the combination of the conserved HBD and the catalytic domain (plus its immediately upstream linker) is needed for the much weaker binding to dsDNA or dsRNA. In contrast, the second HBD weakens binding both to hybrid and to dsDNA (Table 2), consistent with its proposed role in promoting dissociation from the product and facilitating processivity.

**Figure 5.**
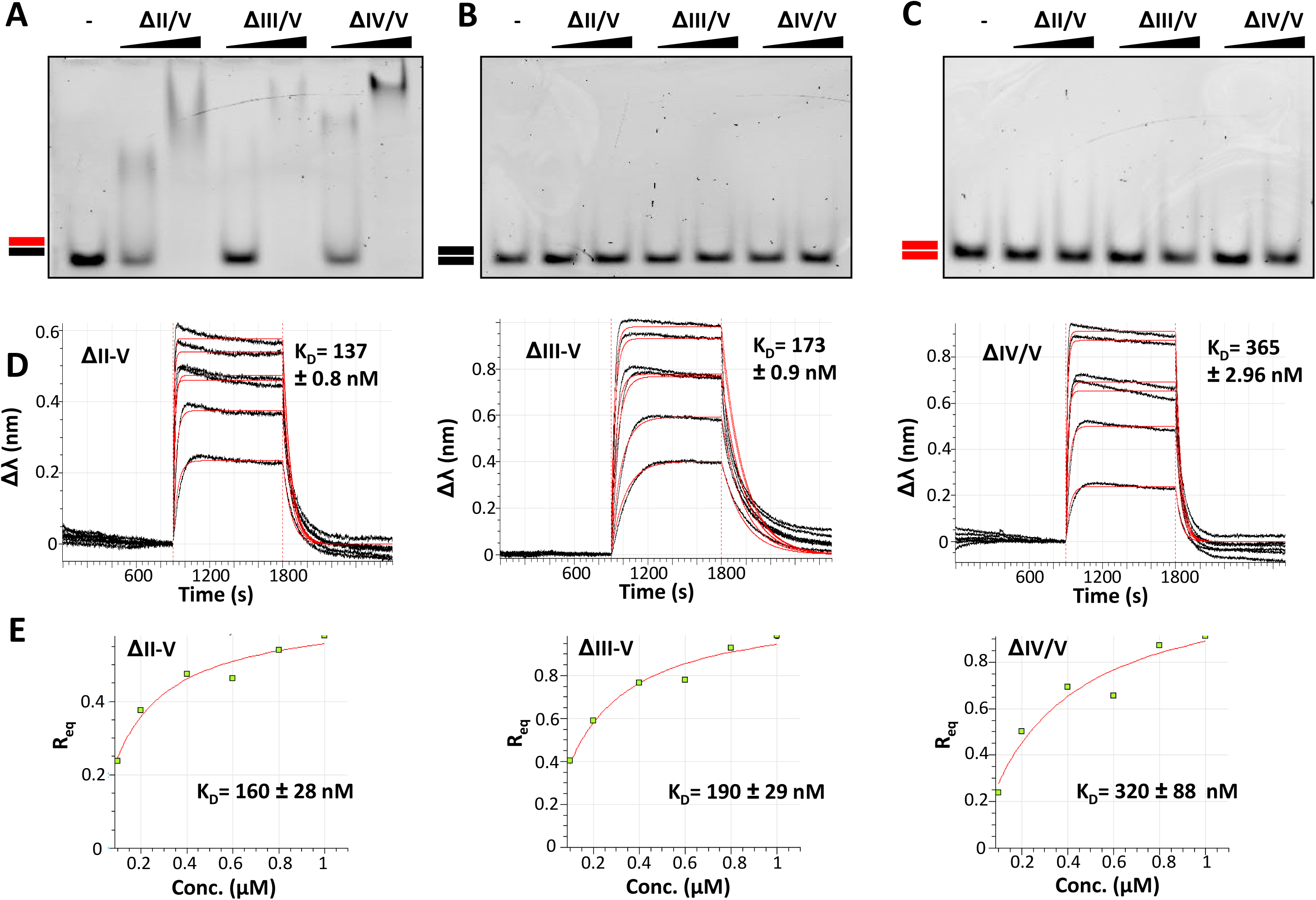
Catalytic domain influences nucleic-acid binding properties of *Dm* RNase H1. (A, B, C) EMSA and (D, E) BLI using *Dm* RNase H1 variants and substrates as indicated, according to the nomenclature of Fig. 2 and 3. The red lines of the sensograms shown in (D) are extrapolated data fitted to a monovalent (1:1) binding model. See Table 3 for association/dissociation parameters. (E) Steady-state analysis of relative equilibrium (R_eq_) plotted against protein concentration. Protein concentrations in (A, B, C) as for Fig. 3, other details as for Fig. 2.

### Protein interactors with RNase H1

The properties of the different regions of *Dm* RNase H1 are strikingly distinct, reflecting the fact that the enzyme must operate in two different cellular compartments and is implicated in a variety of macromolecular processes. To gain further insight into the physiological roles of the protein and its various domains, we initiated a shotgun proteomic screen for proteins that bind to the enzyme. Aiming to identify specific molecular interactors, we applied a whole-cell cross-linking approach (44), initially with just one round of immunopurification, using as bait the full-length protein. As a control to exclude proteins appearing in the list due to non-specific proteotoxic stress provoked by overexpression of a protein targeted to mitochondria, we included mitochondrially targeted YFP (Fig. S7) as well as the previously constructed V5-tagged variants (2) M1V and M16V, with restricted intracellular targeting, respectively to the nucleus and mainly to mitochondria. Finally, with the intent of trapping proteins interacting only transiently with *Dm* RNase H1 during the catalytic cycle, we included also the D252N variant. Raw data for each replicate is shown in Table S2. In each case, we retained target proteins that were identified in every replicate experiment with the given bait protein (n=5 in each case), but which and were absent from all controls (45). The primary screen revealed a list of 63 proteins that we subdivided into two main groups, nuclear (Table 4) and mitochondrial candidates (Table 5), based on the major subcellular location of the protein as currently annotated in Flybase (www.flybase.org). In general, nuclear candidates were found using the full-length, D252N and M1V variants, whereas the mitochondrial candidates were negative using M1V, but few were detected by M16V either, possibly an issue with expression level. The nuclear candidate list included proteins with previously known or inferred roles in heteroduplex processing and DNA replication, whereas the mitochondrial candidates covered a wider spectrum, including many metabolic enzymes not previously implicated in nucleic acid metabolism. Both lists included the respective, compartment-specific single-stranded DNA binding proteins, RpA-70 (as previously reported in mammals; Ref. 6) and mtSSB. Recently, based on *in vitro*-studies of the mammalian proteins, it was proposed that mtSSB and RNase H1 collaborate to generate an RNA primer that would be used by Polγ to initiate mtDNA replication (46). This, and the paucity of other proteins with known or hypothesized roles in DNA transactions amongst the mitochondrial candidates, led us to investigate the interaction between mtSSB and *Dm* RNase H1 in more detail.

**Table 4.** List of nuclear candidates for *Dm* RNase H1 interactors.

| Category | CG number<sup>a</sup> | Official name, gene symbol<sup>a</sup> | Human ortholog<sup>a</sup> | Mean PSM Factor | Detected by which variant(s)? |  |  |  |
| --- | --- | --- | --- | --- | --- | --- | --- | --- |
|  |  |  |  |  | RNase H1 | D252N | M1V | M16V |
| genome maintenance and transcription | CG10279 | Rm62, isoform H, *Rm62* | DDX5, DDX17 | 9 | yes | yes | yes | no |
|  | CG9633 | Replication Protein A 70, *RpA-70* | RPA1 | 6.6 | no | no | yes | no |
|  | CG7831 | Non-claret disjunctional, *ncd* | KIFC1 | 4 | no | no | yes | no |
|  | CG5499 | Histone H2A variant, *His2Av* | H2AFV, H2AFZ | 3 | no | no | yes | no |
|  | CG4747 | Nucleosome-destabilizing factor, *Ndf* | GLYR1 | 2.6 | no | no | yes | no |
|  | CG4206 | Minichromosome maintenance 3, *Mcm3* | MCM3 | 1.8 | no | no | yes | no |
|  | CG7538 | Minichromosome maintenance 2, *Mcm2* | MCM2 | 1 | no | no | yes | no |
| cell cycle progression | CG6392 | CENP-meta, *cmet* | CENPE | 1 | yes | no | no | no |
| other | CG5436 | Heat shock protein 68, *Hsp68* | HSPA1A/HSPA1B | 4.8 | yes | no | yes | yes |

<sup>a</sup>based on current information in Flybase (flybase.org)

**Table 5.** List of mitochondrial candidates for *Dm* RNase H1 interactors.

| Category | CG number<sup>a</sup> | Official name, gene symbol<sup>a</sup> | Human ortholog<sup>a</sup> | Mean PSM Factor | Detected by which variant(s)? |  |  |  |
| --- | --- | --- | --- | --- | --- | --- | --- | --- |
|  |  |  |  |  | RNase H1 | D252N | M1V | M16V |
| metabolism | CG10622 | Succinyl-coenzyme A synthetase $\beta$ subunit, GDP-forming, *ScsβG* | SUCLG2 | 9.6 | yes | yes | no | yes |
|  | CG7010 | Pyruvate dehydrogenase E1 component subunit alpha, *Pdha* | PDHA2 | 7.6 | yes | yes | no | yes |
|  | CG8778 | CG8778 [enoyl-CoA hydratase], *CG8778* | AUH<sup>b</sup> | 6.2 | yes | yes | no | no |
|  | CG7920 | CG7920, isoform A, *CG7920* | [MATN2]<sup>c</sup> | 6 | yes | yes | no | yes |
|  | CG6439 | Isocitrate dehydrogenase 3b, *Idh3b* | IDH3B | 6 | yes | no | no | no |
|  | CG12262 | Medium-chain acyl-CoA dehydrogenase, *Mcad* | ACADM | 5 | yes | no | no | no |
|  | CG4703 | Arc42 [Short-chain acyl-CoA dehydrogenase], *Arc42* | ACADS | 4.8 | yes | yes | no | yes |
|  | CG5889 | Malic enzyme, *Men-b* | ME3 | 4.8 | yes | no | no | no |
|  | CG9006 | Enigma, *Egm* | ACAD9 | 3.8 | yes | yes | no | no |
|  | CG5320 | Glutamate dehydrogenase, *Gdh* | GLUD1 | 3.6 | yes | no | no | no |
|  | CG5599 | CG5599, *CG5599* | DBT<sup>d</sup> | 3.4 | yes | no | no | no |
|  | CG5590 | CG5590, *CG5590* | HSDL2 | 3 | yes | yes | no | no |
|  | CG10639 | L-2-hydroxyglutarate dehydrogenase, *L2HGDH* | L2HGDH | 3 | yes | no | no | no |
|  | CG16935 | CG16935 [Trans-2-enoyl-CoA reductase (NADPH)], *CG16935* | MECR | 2.6 | yes | yes | no | no |
|  | CG1236 | CG1236 [Glyoxylate and hydroxypyruvate reductase], *CG1236* | GRHPR | 2.6 | yes | no | no | no |
|  | CG4860 | CG4860 [Short-chain acyl-CoA dehydrogenase], *CG4860* | ACADS | 2.2 | yes | no | no | no |
|  | CG10194 | CG10194, *CG10194* | NUDT19 | 2.2 | yes | no | no | no |
|  | CG5028 | CG5028, [Isocitrate dehydrogenase (NAD(+))], isoform C, *CG5028* | IDH3G | 2 | yes | yes | no | no |
|  | CG7842 | bad egg, *beg* | MCAT | 2 | yes | no | no | no |
|  | CG12140 | Electron transfer flavoprotein-ubiquinone oxidoreductase, *Etf-QO* | ETFDH | 1.8 | yes | no | no | no |
|  | CG4094 | Fumarase 1, *Fum1* | FH | 1.8 | yes | no | no | no |
|  | CG10672 | CG10672 [Carbonyl reductase (NADPH)], *CG10672* | DHRS4 | 1.6 | yes | no | no | no |
|  | CG3267 | Methylcrotonoyl-CoA carboxylase 2, *Mccc2* | MCCC2 | 1.6 | yes | no | no | no |
| Translation | CG6050 | mitochondrial elongation factor Tu, *mEFTu1* | TUFM | 8.2 | yes | yes | no | yes |
|  | CG6412 | Mitochondrial elongation factor Ts, *mEFTs* | TSFM | 2.4 | yes | yes | no | no |
|  | CG2957 | Mitochondrial ribosomal protein S9, *mRpS9* | MRPS9 | 2.2 | yes | no | no | no |
|  | CG13126 | CG13126, *CG13126* | METTL17 | 2 | yes | no | no | no |
|  | CG5242 | Mitochondrial ribosomal protein L40, *mRpL40* | MRPL40 | 1.6 | yes | no | no | no |
|  | CG5012 | Mitochondrial ribosomal proteinL12, *mRpL12* | MRPL12 | 1.4 | no | yes | no | no |
|  | CG7494 | Mitochondrial ribosomal protein L1, *mRpL1* | MRPL1 | 1.2 | yes | no | no | no |
| protein import and processing | CG11779 | CG11779, *CG11779* | TIMM44 | 8.2 | yes | yes | no | no |
|  | CG8728 | CG8728, *CG8728* | PMPCA | 6.6 | yes | yes | no | no |
|  | CG6155 | Roe1, *Roe1* | GRPEL1 | 3 | yes | yes | no | no |
|  | CG3107 | CG3107, *CG3107* | PITRM1 | 2 | yes | no | no | no |
| OXPHOS | CG2286 | NADH-ubiquinone oxidoreductase 75 kDa subunit, *ND-75* | NDUFS1 | 7.6 | yes | yes | no | no |
|  | CG1970 | NADH dehydrogenase (ubiquinone) 49 kDa subunit, *ND-49* | NDUFS2 | 2.2 | yes | yes | no | no |
|  | CG14724 | Cytochrome c oxidase subunit 5A, *COX5A* | COX5A | 1.4 | yes | no | no | no |
|  | CG10340 | CG10340, *CG10340* | ATPAF1 | 1 | yes | no | no | no |
|  | CG3214 | NADH dehydrogenase (ubiquinone) B17.2 subunit, *ND-B17.2* | NDUFA12 | 1 | yes | no | no | no |
| tRNA metabolism | CG7479 | Leucyl-tRNA synthetase, mitochondrial, *LeuRS-m* | LARS2 | 5.4 | yes | no | no | no |
|  | CG16912 | Tyrosyl-tRNA synthetase, mitochondrial, *TyrRS-m* | YARS2 | 1.2 | yes | no | no | no |
| mtDNA maintenance | CG4337 | Mitochondrial single-stranded DNA-binding protein, *mtSSB* | SSBP1 | 4.2 | yes | yes | no | no |
| oxidative stress | CG5826 | Peroxiredoxin 3, *Prx3* | PRDX3 | 3 | yes | yes | no | no |
|  | CG7217 | Peroxiredoxin 5, *Prx5* | PRDX5 | 3 | yes | yes | no | no |
|  | CG10964 | Sniffer, *sni* | RDH5 | 1 | yes | no | no | no |
| other | CG8479 | Optic atrophy 1 ortholog, isoform D, *Opa1* | OPA1 | 6.4 | yes | no | no | no |
|  | CG14434 | CG14434, *CG14434* | --- | 4 | yes | no | no | no |
|  | CG13850 | CG13850, *CG13850* | TBRG4 | 4 | yes | no | no | no |
|  | CG2794 | CG2794, *CG2794* | --- | 3.4 | yes | no | no | no |
|  | CG11624 | Ubiquitin-63E, *Ubi-p63E* | UBC | 3 | no | yes | no | no |
|  | CG5844 | Spliceosome-ribosome linker protein, *Srlp* | --- | 2.6 | yes | yes | no | no |
|  | CG5915 | Rab7, *Rab7* | RAB7A | 1.8 | yes | no | no | yes |
|  | CG11267 | CG11267 [Hsp10 chaperonin subunit], *CG11267* | HSPE1 | 1.4 | yes | yes | no | no |
|  | CG8993 | CG8993, *CG8993* | TXN2 | 1 | yes | no | no | no |

<sup>a</sup>based on current information in Flybase (flybase.org). Some of these genes are still officially identified only by CG numbers, although orthology indicates a clear enzymatic function also reported in Flybase, and shown here [in brackets].
<sup>b</sup>human ortholog is an RNA-binding variant of the metabolic enzyme
<sup>c</sup>closest match but does not fulfil all criteria for being a true ortholog
<sup>d</sup>human ortholog is dihydrolipoamide branched chain transacylase E2

### mtSSB does not interact directly with RNase H1

Because mass spectrometry revealed mtSSB as a potential interactor with *Dm* RNase H1, we performed co-immunoprecipitation experiments on S2 cells overexpressing V5/His epitope-tagged RNase H1, HA epitope-tagged mtSSB and mtYFP (as control). The subcellular localization of mtSSB-HA and mtYFP were validated by immunocytochemistry (Fig. S7). Western blot analysis revealed that the proteins co-immunopreciptated with RNase H1-V5/His did not include detectable amounts of mtSSB-HA (Fig. 6A and S8, left-hand panels). Similarly, the proteins co-immunopreciptated with mtSSB-HA did not include detectable amounts of RNase H1-V5/His (Fig. 6A and S8, right-hand panels).

**Figure 6.**
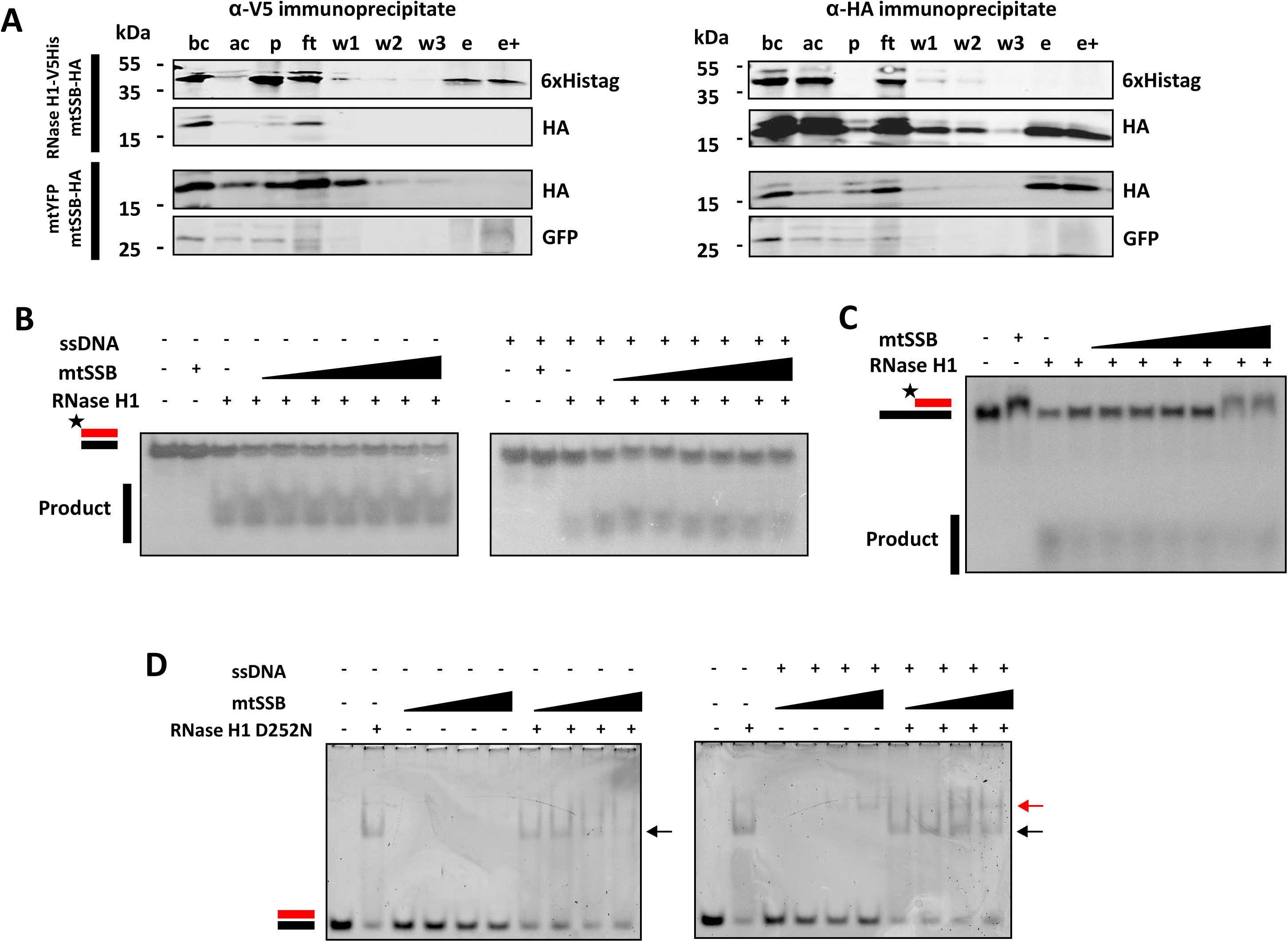
Mitochondrial single strand binding protein does not stimulate RNase H1 activity *in vitro*. (A) Western blots of immunoprecipitates from S2 cells co-expressing RNase H1-V5/His and mtSSB-HA or mtSSB-HA and mtYFP. Immunoprecipitates using anti-V5 (left-hand panels) or anti-HA (right-hand panels) from cells transfected as indicated and probed as shown (to the right of blot panels. All protein extracts were tracked by successive sampling during the procedure, indicated as follows (bc – before crosslinking, ac – after crosslinking, p – pellet, ft – flowthrough, w1, 2 and 3 – washes, e – eluate, still cross-linked, e+ – eluate after reversal of cross-linking). Samples were imaging by Western blot using anti-HA for detecting mtSSB-HA, anti-6x-His tag for detecting RNase H1-V5/His and anti-GFP for detecting YFP. (B, C) Nuclease assays using 0.1 nM of *Dm* RNase H1 and radiolabeled substrates as illustrated according to the nomenclature of Fig. 2, pre-incubated with increasing concentration of mtSSB (0, 5, 10, 25, 50, 100, 250 and 500 nM, without (B, *left panel* and C) or with (B, *right panel*) a 60 nt ssDNA) for 10 min. Substrate in D had a 60 nt ssDNA 3′ overhang. (D) EMSA reactions with 1 µM RNase H1 D252N variant, incubated with increasing concentrations (1, 2.5, 5, 7.5 μM) of free mtSSB (left panel) or ssDNA + mtSSB complex (right panel), for 10 min at room temperature. Arrowheads indicate the complex formed between RNase H1 and the 30 bp RNA/DNA hybrid (black) and between mtSSB and the 60 nt ssDNA (red).

Despite this negative finding with the epitope-tagged proteins *in vivo*, we investigated the functional interactions of the two purified proteins *in vitro*. Addition of mtSSB in large molar excess (minimally 50-fold) had a minor stimulatory effect on the activity of RNase H1 (Fig. 6B), but this was abolished when the assay was conducted in the presence of ssDNA (Fig. 6B). Conversely, when a region of ssDNA was incorporated into the RNase H1 substrate (Fig. 6C), mtSSB appeared to inhibit the RNase H1 reaction slightly, whilst RNase H1 had no effect on complex formation between ssDNA and mtSSB (Fig. 6C, 6D). mtSSB did not affect the formation of complexes between RNA/DNA hybrid and catalytically inactive RNase H1 (Fig. 6D), regardless of the presence of ssDNA.

## DISCUSSION

In this study, we evaluated the biochemical properties of *Dm* RNase H1, determined the functional roles of each region of the protein, and probed for interacting proteins from the two cellular compartments in which RNase H1 is localized. The enzymatic properties of the *Drosophila* enzyme are broadly similar to those of that from humans. However, the two enzymes exhibit different temperature dependencies, but both are highly temperature-sensitive. We identified strong binding for RNA/DNA hybrid in the conserved HBD (region I), and weaker hybrid-binding to the catalytic domain (region V). The HBD also exhibited weak binding for dsDNA and dsRNA, but only in constructs also retaining the catalytic domain. The presence of the intervening domain (III), which we postulated initially as being a second HBD based on structure predictions, had a negative effect on overall hybrid or dsDNA binding affinity, but was also required for full catalytic activity in the presence of the conserved HBD. These properties are summarized in Fig. 7. We identified one major interacting protein from mitochondria, mtSSB. However, studies *in vivo* (Fig. 6A, S8) and *in vitro* (Fig. 6B, 6C, 6D), suggest that the interaction is either transient or indirect, requiring the involvement of other proteins or nucleic acid moieties to mediate or stabilize it.

**Figure 7.**
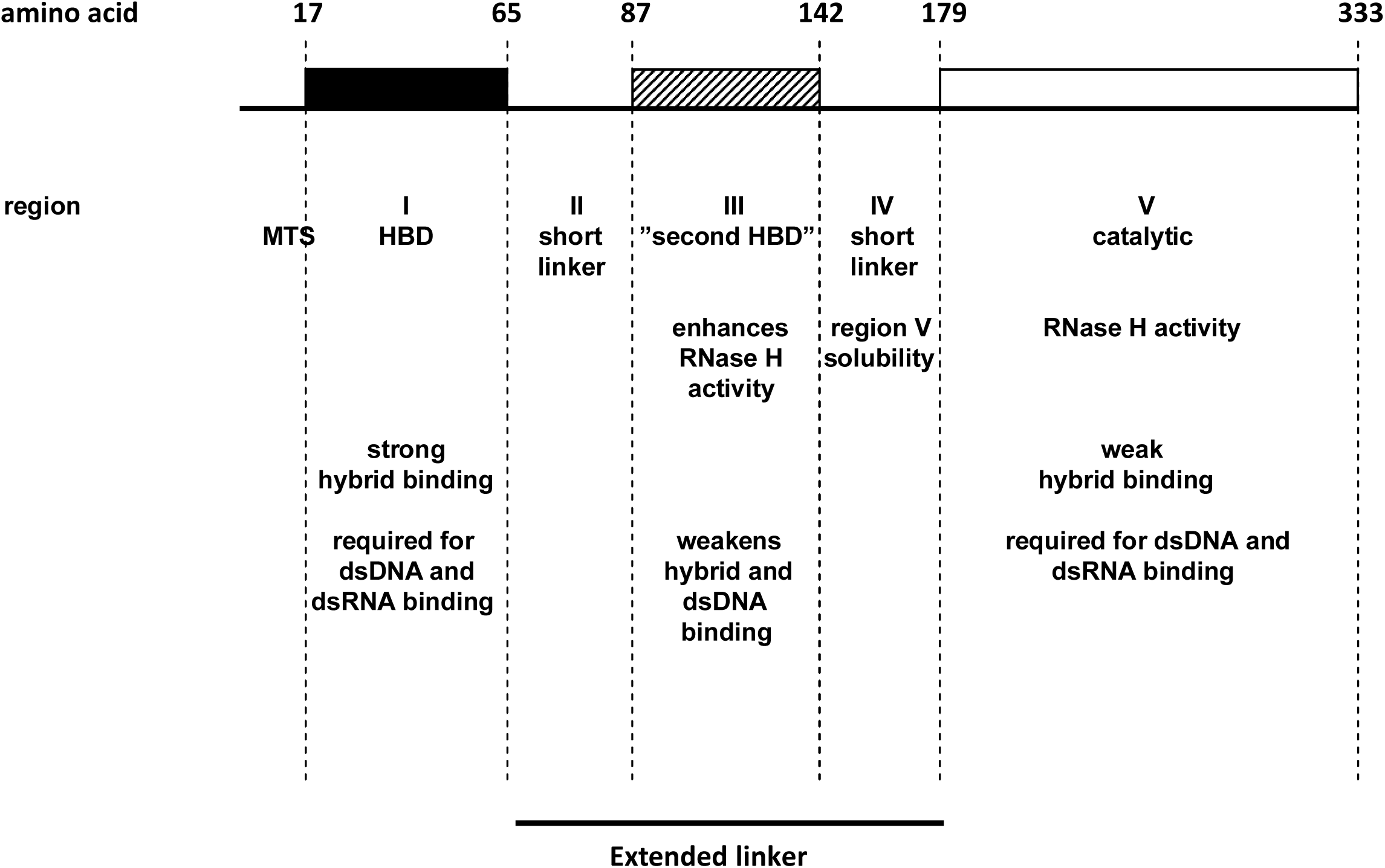
Functional summary diagram of *Dm* RNase H1. Schematic representation of *Drosophila melanogaster* RNase H1. The 5 regions are delimitated as shown, by amino acids 17, 65, 87, 142, 179 and 333. The black box represents the conserved HBD, the hatched box the second HBD and the empty box the catalytic domain. Each region has a short description of its function, based on the experimental findings.

### Functional characterization of the domains of *Dm* RNase H1

Aiming to understand the role of each domain of the enzyme, we created a series of deletion constructs, having divided *Dm* RNase H1 into 5 regions, in order to study the properties conferred by each of them on the enzyme (Fig. 1). Despite the absence of conserved amino acids proposed to be involved in ribonucleotide binding, region I, the conserved HBD, bound RNA/DNA hybrid with a similar affinity as the human HBD (4). The conserved HBD was also required for dsRNA and dsDNA binding (Fig. 4) but, in contrast to the human HBD, it requires the additional presence of the catalytic domain (regions IV and/or V) for these substrates to be bound (Fig. 5, S6). The changes in enzyme kinetics and nucleic acid binding brought about by deletion of the conserved HBD and the adjacent domains suggest that the HBD contributes to RNA cleavage by promoting interaction with heteroduplex. In addition, we observed that the ablation of the HBD abolishes supershifting upon EMSA analysis, suggesting that two protein monomers may associate with a single substrate, conferring processivity to the enzyme, as previously suggested for the mammalian enzyme (5). Regions II and III initially attracted our attention, due to variability in length and composition among eukaryotes. In the *Drosophila* enzyme this extended linker region is particularly long (Fig. 1), with a predicted isoelectric point (pI, 5.18) similar to that of mammals or zebrafish, but lower than in *Xenopus tropicalis* (9.11) or *Caenorhabditis elegans* (6.79). Moreover, structural modeling suggested that region III may constitute a second HBD (Fig. S1C). Although we were unable to study its properties in isolation, due to its insolubility, our other data (Tables 2, 3) indicate that, whereas this region actually weakens the overall hybrid-binding of the enzyme (Fig. 3), it enhances catalysis (Table 1). Although a conclusive interpretation of these findings must await full elucidation of the reaction mechanism of *Dm* RNase H1, it is tempting to suggest that the HBD-like fold of domain III is involved in shuttling substrate from the conserved HBD to the catalytic domain, as part of the processivity mechanism. Its deletion would thus promote tight and persistent hybrid binding by the conserved HBD, leading to a lower catalytic throughput. An alternative explanation for the findings might be that the length of the extended linker *per se* determines the catalytic activity and binding affinity of the enzyme. In other words, a long linker allows the catalytic domain to interact processively or successively with substrate, whilst also influencing binding (a ‘running dog on a leash’ model). Whether the ‘second HBD’ actually binds nucleic acid, even transiently, and whether its predicted fold is functionally important, must await detailed structural analysis and further mutagenic studies of the enzyme, using crystallography. The differences in the binding properties of the enzyme *in vitro*, using the linear hybrid (Fig. 2) and R-loop substrates (Fig. 3), may reflect functionally important differences *in vivo*, such as in the removal of persistent heteroduplex regions that arise during transcription, *versus* the processing of DNA replication intermediates.

The catalytic domain was predicted to adopt a similar fold as previously observed in viruses (47), bacteria (7, 8, 48, 49) and mammals (9), and includes the conserved DEDD motif (Fig. S1), from which residue D252 was shown to be essential for activity, as in other organisms (16). Importantly, the catalytic domain was required for the 2:1 binding model (Table 3), but not for the supershift seen in EMSA (Fig. 5A), for which the conserved HBD alone was sufficient. This suggests that the reaction involves not only the binding of a second protein molecule but also, potentially, a conformational change dependent on the catalytic domain, which may enable the second protein moiety to bind.

### Functional comparison with RNase H1 from other species

Most studies of eukaryotic RNase H1 have focused on the enzymes from yeast and from humans, whilst little was known about the function, role and biochemical properties of the *Drosophila* enzyme (2, 9, 36). We initially characterized the enzymatic activity of *Dm* RNase H1 at 30 °C, revealing slightly different kinetic parameters from those of the human enzyme, studied previously at 37 °C (16, 28). 30 °C has widely been used as a reference temperature for studies of *Drosophila* enzymes, notably those involved in mtDNA metabolism. It represents a temperature about 10-12 °C warmer than the typical physiological temperature of the fly. However, assuming that, like their mammalian counterparts, *Drosophila* mitochondria are 10-12 °C warmer than the cells and tissues in which they function (50), 30 °C should represent an optimal temperature at which to study mitochondrially localized enzymes. When we compared the human and *Drosophila* enzymes we found their properties to be highly influenced by temperature, with the human enzyme essentially inactive at 30 °C, but the *Drosophila* enzyme exhibiting a much lower substrate affinity (higher K_M_) at 37 °C than at 30 °C. At its presumed optimal temperature of 30 °C, the *Drosophila* enzyme displayed a markedly higher affinity and catalytic turnover rate than the human enzyme at 37 °C (Table 1). However, given the marked temperature sensitivity of both enzymes, and the fact that the *in vivo* operating temperature of the human enzyme in mitochondria is probably much closer to 50 °C than to 37 °C (50), the two enzymes may have more similar properties than is apparent. Note also that the cell nucleus should be much closer to ambient temperature (in the fly) or to 37 °C in humans, such that the enzymatic properties of RNase H1 in the nucleus may differ substantially.

Similar caution should apply to measurements of affinity constants, especially given the uncertainties raised by the use of different methods in the various studies. Here, applying BLI using the D252N-substituted enzyme, we inferred a K_D_ value intermediate between those previously reported for human (16) and *E. coli* RNase H1 (51). Previously, BLI has been used to measure nucleic acid-protein interactions, obtaining similar values as with other approaches (52), and has been specifically applied to the study of Polγ (53).

Mammalian, yeast and *E. coli* RNase H1 bind dsRNA, but only the enzyme from the archaeon *Sulfolobus tokodaii 7* has been demonstrated to digest this substrate (54). In the present study we found that *Dm* RNase H1 also binds dsRNA and dsDNA (Fig. 4, Table 2), respectively ∼10-fold and ∼100-fold less tightly than RNA/DNA hybrid, but does not digest these substrates, nor does it bind or digest ssRNA or ssDNA, properties shared with the human and *E. coli* enzymes (55).

In a previous study, RNase H1 from the budding yeast *Saccharomyces cerevisiae* was found to have two nucleic acid-binding domains located in the N-terminal region (41; see Fig. 1),with the more N-terminally located such region showing a much higher affinity for substrate. This raises the question as to whether the architecture and suggested reaction mechanism of the *Dm* enzyme, with two HBDs, might also apply in yeast. However, in other respects the *Dm* enzyme differs fundamentally from that of *S. cerevisiae*, having much lower affinity for dsRNA than for RNA/DNA hybrid, whilst the ‘HBD’ of the yeast enzyme actually binds dsRNA more tightly than hybrid (41). Furthermore the catalytic domain of the *Dm* enzyme binds hybrid on its own, and is required for binding to dsRNA, whilst the enzyme from *S. cerevisiae* shares neither property (41). Thus, the functional properties of the two enzymes *in vivo* are likely to differ substantially.

### Significance of RNase H1 interactors

RNase H1 has been implicated in several processes in different sub-cellular compartments. Our mass spectrometry analysis revealed a list of potential interactors in the nucleus, as well as in mitochondria. In mammals, RPA (replication protein A) has been shown to recruit RNase H1 to R-loops and stimulate its enzymatic activity (6), via an interaction with the HBD. Such an interaction facilitates the role of the enzyme in heteroduplex surveillance and also, potentially, in DNA maintenance. SSB and RNase H1 have also been reported to interact in *E. coli* (56), in this case via the catalytic core, because the bacterial enzyme lacks the HBD. RPA (RpA-70) was one of the top nuclear hits in our screen for interacting proteins (Table 4), which also yielded two subunits of the nuclear replicative helicase, Mcm2 and Mcm3 (57, 58), whilst RPA has been implicated in nuclear processes other than DNA replication (59, 60). We also identified mtSSB, the functional homolog of RPA in mitochondria, as a prominent hit (Table 5). mtSSB is a well characterized component of the mtDNA replication machinery (61), whilst RNase H1 in *Drosophila* has also been inferred to play a role in mtDNA replication fork progression (2). Furthermore, mtSSB and RNase H1 have been proposed to co-operate in the initiation of mtDNA replication (46), which spurred us to examine the possible interaction between them in more detail. The two proteins did not co-immunoprecipitate when overexpressed together *in vivo* (Fig. 6A, S8), nor did enzymatic assays and EMSA reveal convincing evidence for any direct interaction *in vitro* (Fig. 6B, 6C, 6D), apart from a very slight stimulation or inhibition of the enzyme, depending on the specific substrate (Fig. 6B, 6C). Although overexpression *in vivo* and *in vitro* analysis of the properties of bacterially-expressed proteins are subject to different potential artifacts, the fact that neither approach supported a direct interaction implies that mtSSB and *Dm* RNase H1 most likely interact indirectly *in vivo*, requiring an unidentified partner protein or nucleic acid moiety to have enabled their co-detection by mass spectrometry. This does not exclude that the two proteins may co-operate as suggested (46), with mtSSB binding to the single-stranded DNA displaced at an R-loop, promoting RNase H1 recruitment that would partially digest the annealed RNA, thus creating a 3′ end accessible for extension by Polγ. However, it would imply that at least one additional partner would be required for such a recruitment to occur. This partner cannot simply be ssDNA, because it did not facilitate any direct interaction between the proteins *in vitro* (Fig. 6B, 6C, 6D). It is also possible that mtSSB and RNase H1 co-localize at the replication origin, at replication forks and at dispersed R-loops only by virtue of their substrates (ssDNA and RNA/DNA hybrid) being juxtaposed at these sites, and thus that their close association is purely adventitious. Importantly, negative findings such as ours, even though supported by multiple approaches, may be erroneous if the conditions for analysis *in vitro* are inappropriate. Future experiments using different methods may be needed to confirm (or revise) the apparent absence of direct interaction. The functional interactions of mtSSB with other mitochondrial replication proteins, such as Polγ (62) or mtDNA-helicase (63) require low salt conditions, similar to those used here, which yielded negative findings. However, it is possible that some other feature of the intramitochondrial environment is required to maintain mtSSB/RNase H1 links in *Drosophila*.

In human cells, mtSSB has been reported to localize partially to RNA granules (64), whilst defects in the machinery of RNA processing and degradation results in the accumulation of persistent R-loops, resulting in RNase H1 recruitment to nucleoids (65). In an earlier study, vertebrate RNase H1 was not observed as a nucleoid protein (66), consistent with its interactions with mitochondrial replication and RNA processing enzymes being transient, and mediated by its association with RNA/DNA hybrid substrate, rather than by direct protein-protein interactions

Amongst other nuclear hits, we identified a second component of the R-loop processing machinery, Rm62, the *Drosophila* homologue of human DDX5 (67). This suggests a potential interaction between two independent machineries to resolve R-loops. The list of mitochondrial candidates was much longer, and included, as a prominent class, many metabolic enzymes involved in core processes such as fatty acid oxidation, the TCA cycle and OXPHOS, as well as some proteins involved in mitochondrial translation. Metabolic enzymes have been previously reported as at least peripheral components of nucleoids in many species, and there is ongoing debate as to whether this association is meaningful. In regard to the present study, the question arises as to whether they represent ‘real’ interactors with RNase H1 or are just passenger proteins brought along by cross-linking in a protein-rich environment. Some hits, such as the fly orthologs of human TIMM44, GRPEL1, PITRM1 and PMPCA, are likely to be artifacts of overexpression, resulting from the machinery of protein import and processing becoming overwhelmed, even though these proteins did not appear in the mtYFP negative control list. Given the fact that the mitochondrial candidate list is ‘over-inclusive’ in this manner, and that many nuclear hits are congruent with previous data or with assumptions made on the basis of such data, the absence of any known component of the apparatus of mitochondrial nucleic acid metabolism other than mtSSB is striking.

Note that a number of other possible candidates do not figure in Tables 4 and 5 because of the strict exclusion criteria, i.e. they were found in at least one control replicate (see Table S2). Prominent amongst these was P32 (CG6459 in *Drosophila*), reported as being present in both the mitochondria and nucleus and implicated in diverse processes, and which was previously found to associate with RNase H1 in mammals and proposed to stimulate its activity (28). Whilst it may be enriched in the fraction associating with RNase H1, it is not specific to this fraction.

Having undertaken this study to follow up previous observations that a deficiency of *Dm* RNase H1 results in characteristic abnormalities of mtDNA replication (2), reflecting aspects of the transcriptional map of *Drosophila* mtDNA, the absence of mitochondrial hits for mitochondrial replication and transcription proteins other than mtSSB was unexpected. Furthermore, the clinical features manifested by patients with mutations in the *RNASEH1* gene resembles those associated with other disorders caused by defects in the mtDNA replication apparatus (21-23). One simple explanation is that RNase H1 does not interact directly with replication proteins, and that the effects of its deficiency on mtDNA replication are secondary to a failure to process R-loops and other hybrid-containing structures. In other words, RNase H1 may function independently and not be part of the mitochondrial replisome, transcriptional machinery or any other protein complex within or associated with the nucleoid.

## EXPERIMENTAL PROCEDURES

### Molecular modeling

*Dm* RNase H1 protein structure was modeled by SWISS-MODEL (Biozentrum, University of Basel, Switzerland; 38-40). Its amino acid sequence was compared to those of proteins from the protein data bank, PDB, with crystallographically-determined structures, to find potential templates, ranking them based on estimated GMQE (Global Model of Quality Estimation). Model quality was assessed by QMEAN, a value calculated by global and local geometrical characteristics of the model with respect to the template. Models were analyzed and visualized by PyMol (Schrödinger).

### Cloning into expression vectors

*rnh1* and *mtSSB* cDNAs were derived by PCR using methods described previously (2) and primers as listed in Table S1. The *rnh1* cDNA was cloned into pET-26b(+) (Novagen, Merck Millipore) for bacterial expression with an in-frame C-terminal 6xHis tag. Partially deleted *rnh1* variants (Fig. 1B) were generated by a two-step PCR procedure, as described previously (2). PCR-based site-directed mutagenesis was used to create variant cDNAs bearing the D252N point mutation, predicted to abolish nuclease activity in RNase H1. For expression in *Drosophila* S2 cells (68) the *mtSSB* coding sequence was cloned into pMT-puro (Addgene) using a two-step procedure. It was first amplified and inserted into pMT/V5-His B (ThermoFisher Scientific), using primers that introduced EcoRI and XhoI restriction sites, then recloned into pMT-puro using primers that added KpnI and PmeI restriction sites, also eliminating the V5-His tag but adding a C-terminal HA tag in its place (see Table S1). Successfully transfected colonies were selected by plating on 0.5 μg/ml puromycin (InvivoGen). All plasmids were sequenced before use.

### Protein expression and purification

Competent BL21 Star™ (DE3) *E. coli* cells (ThermoFisher Scientific) were transformed with pET-26b(+)-derived DNA constructs using heat shock, and selected on 50 μg/ml kanamycin plates. Cells from single colonies were grown overnight in 200 ml L-broth (LB, 1% tryptone, 1% NaCl, 0.5% yeast extract, all w/v), supplemented with 50 μg/ml kanamycin, at 37 °C with shaking at 250 rpm. The culture was then diluted into 4 l of LB (in eight two-liter Erlenmeyer flasks) and incubated at 37 °C with shaking until it reached an OD_600_ of ∼0.7. Protein expression was induced by the addition of IPTG (isopropyl β-D-1-thiogalactopyranoside) to 400 µM and a further incubation for 3 h. Cells were harvested by 10 min centrifugation at 5,000 *g*_*max*_ and stored at - 80 C°. Bacterial pellets were thawed in ice-cold resuspension buffer (30 mM Tris-HCl, 200 mM KCl, 2 mM DTT, 10% glycerol, pH 8.0) containing, per 25 ml, one cOmplete™ EDTA-free Protease Inhibitor Cocktail tablet (Roche). Lysozyme (Invitrogen) was added to 200 μg/ml on ice for 40 min, after which cells were lysed by three rounds of sonication on ice, using a Vibra Cell (VC) 505 sonicator (Sonic & Materials, Inc.), fitted with a 13 mm probe, set to 60% amplitude with a 1 s/2 s on/off cycle for 3 min. Where needed, a fourth round of sonication was added. The lysate was ultracentrifuged at 100,000 *g*_*max*_ for 30 min at 4 °C, after which the supernatant was filtered through a 0.45 µm nylon syringe-filter (GE Healthcare Whatman™ Uniflo) and loaded dropwise overnight onto a 3 ml Ni-NTA agarose (ThermoFisher Scientific) column (Qiagen 30230) pre-washed in water then pre-equilibrated with equilibration buffer (30 mM Tris-HCl, 200 mM KCl, 2 mM DTT, 10% glycerol, 5 mM imidazole, pH 8.0, plus the protease inhibitor). All chromatography and gel filtration steps were conducted at 4 °C. Non-specifically bound proteins were removed by successively washing with 15 ml buffer 1 (30 mM Tris-HCl, 200 mM KCl, 2 mM DTT, 10% glycerol, 25mM imidazole, pH 8.0, plus the protease inhibitor) and 15 ml buffer 2 (30 mM Tris-HCl, 200 mM KCl, 2 mM DTT, 10% Glycerol, 50 mM imidazole, pH 8.0, with same protease inhibitor), after which the desired protein was finally eluted with 6 ml buffer 3 (30 mM Tris-HCl, 1 M KCl, 2 mM DTT, 10% glycerol, 250 mM imidazole, pH 8.0, plus the protease inhibitor) and collected in 0.5 ml aliquots. Fractions containing the desired protein were pooled and gel filtration was performed on Superdex 75 or 200 10/300 GL columns (GE Healthcare) mounted into an ÄKTA P100 chromatography system (GE Healthcare). Columns were washed and equilibrated with SE buffer (30 mM Tris-HCl, 200 mM KCl, 2 mM DTT, 10% glycerol, pH 8.0) and 0.5 ml fractions were collected after sample injection. Protein purity was analyzed by SDS-PAGE on 12.5% or 15% polyacrylamide gels, followed by Coomassie-Blue staining. The purified proteins were aliquoted and stored at −80 °C. *Drosophila* mtSSB was expressed and purified as described previously (69).

### Nucleic acid substrates

RNA and DNA oligonucleotides used for the experiments are listed in Table S1. For testing nuclease activity, DNA or RNA oligonucleotides were 5′ radiolabeled with [γ-^32^P]ATP (PerkinElmer, 3000 Ci/mmol), using T4 Polynucleotide Kinase (New England Biolabs) as described in manufacturer’s protocol. The radiolabeled nucleic acid was recovered by gel-filtration using a Sephadex G-50 fine Quick Spin column (Roche) according to manufacturer’s instructions, and its concentration was estimated by scintillation counting. To generate a double-stranded substrate, a two-fold excess of the complementary strand was added and incubated for 5 min in annealing buffer (90 mM Tris-HCl, 10 mM MgCl_2_, 50 mM KCl, pH 8.0) at 95 °C, then bench-cooled to room temperature.

### Nuclease activity assay

Radiolabeled substrate was incubated with purified *Dm* RNase H1, variants or control enzymes as described below and in figure legends, and at the indicated temperatures, in reaction buffer (50 mM Tris-HCl, 2 mM DTT, 5 mM MgCl_2_, 400 μg/ml bovine serum albumin (BSA), pH 8.0) with salt concentration adjusted to 25 mM KCl. Positive control enzymes (ThermoFisher Scientific),were varied according the substrate: for linear RNA/DNA hybrids and R-loops, RNase H, for dsRNA and ssRNA, RNase A and for dsDNA and ssDNA, DNase I. Reactions were stopped with 10x STOP solution (10% SDS (w/v), 100 mM EDTA), incubated for 10 min at 60 °C, separated by non-denaturing polyacrylamide gel (1x TBE, 12.5% acrylamide:bis-acrylamide 19:1) electrophoresis and heat/vacuum dried for autoradiography (Amersham Hyperfilm ECL, GE Healthcare). Images were analyzed using ImageJ. For calculating turnover kinetics, initial cleavage rates (V_0_) were calculated for each RNase H1 variant at 0.2 nM, in the presence of different concentrations of radiolabeled hybrid (2.5, 5, 7.5 and 10 nM) at different time points. Product generation was plotted as a function of time and V_0_ was calculated by estimating the time required to generate 10% of the total product. V_0_ was plotted against substrate concentration and fitted to the Michaelis-Menten equation. Kinetic constants were estimated by the Lineweaver-Burk equation.

### Electrophorestic mobility shift assay (EMSA)

Binding reactions were conducted in binding buffer (25 mM Tris-HCl, 1 mM DTT, 10 mM EDTA, 20 μg/ml BSA, pH 8.0) and salt was adjusted to a final concentration of 25 mM KCl. Protein and nucleic acid concentrations are as indicated in figure legends. Samples were incubated at room temperature for 30 min and fractionated by 6% polyacrylamide gel (0.5x TBE, 6% acrylamide: bis-acrylamide 29:1, 2.5% glycerol) electrophoresis in TBE buffer. Non-radiolabeled nucleic acid was stained with GelRED™ (Biotium) for 20 min in TBE and washed for 20 min with TBE. Gel images were analyzed with ImageJ.

### Biolayer interferometry (BLI)

5′-biotinylated (RNA or DNA) oligonucleotides (Sigma, Table S1) were diluted with non-biotinylated complementary oligonucleotides in PBS, each at a concentration of 10 µM. The oligonucleotide mixture (dsDNA, dsRNA or RNA/DNA hybrid) was incubated at 95 °C for 5 min and left to anneal at room temperature overnight. Streptavidin-coated biosensors (FortéBio) were humidified for 30 min in water. This and all subsequent steps were conducted in 384-well plates using 80 μl of solution per sensor. Sensors were transferred to 25 nM annealed, biotinylated nucleic acid solution for 5 min, followed by a quenching step with 10 µg/ml Biocytin (Sigma) diluted in PBS. Sensors were blocked and equilibrated with Kinetics Buffer (30 mM Tris-HCl, 100 mM KCl, 10% glycerol, 10 mM EDTA, 2 mM DTT, 400 µg/ml BSA, pH 8.0) twice for 10 min. Sensors were transferred to Kinetics Buffer containing different protein concentrations for 15 min. Dissociation was measured using an Octet® RED384 BLI detection system (FortéBio), by transferring the sensors to kinetic buffer for 15 min. All steps were conducted at 30 °C with mixing at 1,000 rpm. Processing and analysis of the data were as described (53), using Octet® Systems software (FortéBio). Monovalent (1:1) or heterogeneous (2:1) binding models were used for estimating equilibrium dissociation constant (K_D_), association rate (K_on_) and dissociation rate (K_off_), as indicated in figures and tables for each variant.

### Immunohistochemistry

S2 cell transfection, fixation, staining and imaging were as described previously (2). mtYFP and mtSSB-HA were detected using mouse monoclonal antibodies, respectively against GFP (Abcam ab1218, 1:10,000), and HA tag (2-2.2.14; ThermoFisher Scientific 26183, 1:10,000), used with goat anti-mouse IgG AlexaFluor®568 (ThermoFisher Scientific A-11004, 1:10,000) as secondary antibody. Mitochondrial Cox4 was counter-stained with rabbit polyclonal anti-COXIV antibody (Abcam ab16056, 1:10,000), used with goat anti-rabbit IgG AlexaFluor®488 (ThermoFisher Scientific A-11008, 1:10,000) as secondary antibody. Images were optimized for brightness and contrast using ImageJ but not manipulated in any other way.

### Immunoprecipitation from S2 cells

Stably-transformed S2 cell-derived cell-lines were generated (2) and maintained (68), and protein expression induced as previously (2). Immunopreciptation was conducted essentially as described previously (44). Approximately, 6 × 10^8^ cells were cross-linked with 1% formaldehyde for 10 min at room temperature with continuous agitation. Cross-linking was stopped by adding 125 mM glycine. Cells were harvested by centrifugation, washed four times with ice-cold Tris-buffered saline (TBS), resuspended in RIPA lysis buffer (50 mM Tris-HCl, 150 mM NaCl, 1% v/v Nonidet P40; 0.5% w/v sodium deoxycholate; 0.1% w/v SDS) and incubated for 30 min on ice. Cells were water-bath sonicated (FinnSonic M0, ultrasonic power 200 W, ultrasonic frequency 40 kHz) at 4 °C for 20 min. Cell extracts were incubated at 37 °C for 30 min following the addition of RNase A (ThermoFisher Scientific, 100 μg/ml), DNase I (ThermoFisher Scientific, 5 U/ml), Benzonase® nuclease (Sigma, 50 U/mL), MgCl_2_ to 2.5 mM and CaCl_2_ to 1 mM. Samples were centrifuged for 10 min at 4 °C, after which supernatants were incubated with 3 μl anti-V5 (Invitrogen R-960-25) or anti-HA (Invitrogen 26183) antibody overnight at 4 °C on a rocking shaker. Protein-antibody complexes were collected by incubating protein extracts with 20 μl SureBeads Protein G magnetic beads (Bio-Rad) for 30 min at room temperature on a rocking shaker, followed by three washes with RIPA buffer and resuspension of magnetic beads in protein-loading buffer (PLB: 2% w/v SDS; 2 mM β-mercaptoethanol, 4% v/v glycerol, 40 mM Tris-HCl, 0.01% w/v bromophenol blue, pH 6.8). Crosslinking was reversed by heating at 95 °C for 30 minutes. Samples were analyzed by mass-spectrometry, as below, or by Western blotting, essentially as described previously (2), using the following primary/secondary antibodies: rabbit anti-6x-His Tag (ThermoFisher Scientific PA1-983B, 1:10,000), mouse monoclonal anti-GFP (Abcam ab1218, 1:10,000, used to detect YFP), and mouse monoclonal anti-HA Tag (ThermoFisher Scientific 26183, 1:10,000), used with either IRDve® 680LT anti-mouse (LI-COR, 925-68022, 1:10,000) or IRDve® 680LT anti-rabbit (LI-COR, 925-68023, 1:10,000), as appropriate. Blot images were cropped and rotated for presentation, and optimized for contrast and brightness, but not subjected to other manipulations.

### LC-MS/MS

Liquid chromatography coupled to tandem mass spectrometry (LC-MS/MS) was carried out on an EASY-nLC1000 chromatograph connected to a Velos Pro-Orbitrap Elite hybrid mass spectrometer with nano electrospray ion source (all instruments from ThermoFisher Scientific). The LC-MS/MS samples were separated using a two-column setup, consisting of a 2 cm C18 Pepmap column (#164946, ThermoFisher Scientific), followed by a 15 cm C18 Pepmap analytical column (#164940 ThermoFisher Scientific). Samples were loaded in buffer A and the linear separation gradient consisted of 5% buffer B for 5 min, 35% buffer B for 60 min, 80% buffer B for 5 min and 100% buffer B for 10 min at a flow rate of 0.3 μl/min (buffer A: 0.1% trifluoroacetic acid in 1% acetonitrile; buffer B: 0.1% trifluoroacetic acid in 98% acetonitrile). 6 μl of sample was injected per LC-MS/MS run and analyzed. Full MS scans were acquired with a resolution of 60,000 at 300-1700 m/z range in the Orbitrap analyzer. The method was set to fragment the 20 most intense precursor ions with CID (energy 35). Data was acquired using LTQ Tune software. Acquired MS2 scans were searched against the *Drosophila melanogaster* protein database using the Sequest search algorithms in Thermo Proteome Discoverer. Allowed mass error for the precursor ions was 15 ppm, and for the fragment 0.8 Da. A static residue modification parameter was set for carbamidomethyl +57,021 Da (C) of cysteine residue. Methionine oxidation was set as dynamic modification +15,995 Da (M). Only full tryptic peptides were allowed, with a maximum of 1 missed cleavage.

## ACKNOWLEDGEMENTS

We thank Robert Crouch for supplying purified human RNase H1. We also thank Ian Holt and Maria Solá for advice and useful discussions, Juha Määttä for guidance with BLI and Tea Tuomela, Eveliina Teeri and Merja Jokela for technical assistance.

## FUNDING

This work was supported by Academy of Finland [Centre of Excellence grant 272376 and Academy Professorship grant 256615 to HTJ and Finland Distinguished Professorship to LSK]; The Finnish Cultural Foundation [grant 00190260 to JMGdeC.]; Tampere University; and the Sigrid Juselius Foundation. The work used core facilities for proteomics in the Institute of Biotechnology (Helsinki), as well as for imaging and protein analysis in Tampere, both part-supported by Biocenter Finland.

## CONFLICT OF INTEREST

The authors declare that they have no conflicts of interest with the contents of this article.

